# Multimodal frequency representations are embedded in modality-defined cortical sensory systems

**DOI:** 10.1101/628305

**Authors:** Shoaibur Rahman, Kelly Anne Barnes, Lexi E. Crommett, Mark Tommerdahl, Jeffrey M. Yau

**Author notes:** Corresponding author: Jeffrey M. Yau, One Baylor Plaza, T111, Houston, TX 77030.

## Abstract

Sensory information is represented and elaborated in hierarchical cortical systems that are thought to be dedicated to individual sensory modalities. This traditional view of sensory cortex organization has been challenged by recent evidence of multimodal responses in primary and association sensory areas. Although it is indisputable that sensory areas respond to multiple modalities, it remains unclear whether these multimodal responses reflect selective information processing for particular stimulus features. Here, we used fMRI adaptation to identify brain regions that are sensitive to the temporal frequency information contained in auditory, tactile, and audiotactile stimulus sequences. A number of brain regions distributed over the parietal and temporal lobes exhibited frequency-selective temporal response modulation for both auditory and tactile stimulus events, as indexed by repetition suppression effects. A smaller set of regions responded to crossmodal adaptation sequences in a frequency-dependent manner. Despite an extensive overlap of multimodal frequency-selective responses across the parietal and temporal lobes, representational similarity analysis revealed a cortical “regional landscape” that clearly reflected distinct somatosensory and auditory processing systems that converged on modality-invariant areas. These structured relationships between brain regions were also evident in spontaneous signal fluctuation patterns measured at rest. Our results reveal that multimodal processing in human cortex can be feature-specific and that multimodal frequency representations are embedded in the intrinsically hierarchical organization of cortical sensory systems.

**Significance Statement:** A hallmark of traditional brain organization models is the segregation of signals from the different senses in modality-dedicated brain regions. Recent evidence showing multimodal activity in brain regions thought to be dedicated to a single modality have challenged the traditional sensory cortex model. Notably, few studies have explored the feature-specificity of multimodal responses found in sensory cortex. Here, we used fMRI adaptation to identify parietal and temporal cortex regions which exhibited sensitivity to both tactile and auditory frequency information. These univariate results demonstrate that multimodal processing in sensory cortex can be feature-specific. Using the same data, though, we found clear evidence of modality-based cortical organization estimated from multivariate response patterns and spontaneous BOLD signal fluctuations. Thus, our results reveal an embedding of feature-specific multimodal processing in traditionally-defined cortical systems.

## Introduction

How the brain is organized to support perception of unisensory and multisensory signals is a fundamental question. Traditional organization models posit an initial processing of information in unisensory cortical systems before subsequent processing in multimodal systems residing in association cortices. Yet, it is now incontrovertible that brain regions traditionally considered to be dedicated to a single modality can be modulated or driven by sensory inputs delivered via multiple sensory modalities (Ghazanfar and Schroeder, 2006; Driver and Noesselt, 2008). These discrepancies motivate a debate between the traditional “modality-based scheme” for sensory cortex organization and a “function-based scheme” in which sensory regions perform particular operations regardless of input modality (Pascual-Leone and Hamilton, 2001; Driver and Noesselt, 2008). Here we characterized BOLD fMRI responses to auditory and tactile stimulus sequences to investigate how sensory cortex is organized to support multimodal temporal frequency processing.

In humans, the relative sensitivities of the auditory (0.02-20kHz) and somatosensory (2-1000Hz) systems to temporal frequency information imply that the two senses overlap in the processing of signals below 1000Hz. Environmental cues in this frequency range may be particularly relevant for signaling our interactions with objects (Miller et al., 2018), fine texture information (Yau et al., 2009a; Manfredi et al., 2014), and speech (Titze, 1994; Lattner et al., 2005). There are clear correspondences between auditory and tactile frequency perception (Bensmaia et al., 2005) as well as extensive evidence for frequency-specific audiotactile interactions with flutter (below 60 Hz) (Convento et al., 2019) and supra-flutter signals (>85Hz) (Yau et al., 2009b, 2010, Crommett et al., 2017, 2018). The intimate relationship between auditory and tactile frequency perception presumably reflects the existence of shared or interactive neural representations.

The neural codes for low-frequency auditory (Bendor and Wang, 2007) and tactile stimulation (Romo and Salinas, 2003) in the flutter range are well established and neural populations that analogously signal flutter in both modalities have been identified (Vergara et al., 2016). In contrast, while neural populations are clearly tuned for supra-flutter auditory frequency (Bendor and Wang, 2005; Wang and Walker, 2012), an explicit coding of vibration frequencies – which can be represented in spike timing (Harvey et al., 2013) – has not been established. Accordingly, the relationship between auditory and somatosensory supra-flutter representations remains unclear (Saal et al., 2016). Consistent with a feature-based organization scheme in which auditory cortex is specialized for frequency processing irrespective of input modality, some auditory areas respond to tactile stimulation alone (Foxe et al., 2002; Fu et al., 2003; Kayser et al., 2005; Schurmann et al., 2006). However, other studies (Liang et al., 2013; Perez-Bellido et al., 2017) have revealed that auditory information can also be decoded from putative somatosensory areas. Thus, questions remain regarding the nature of auditory and tactile frequency representations in the human brain and the organization of the cortical systems supporting these representations.

We used an fMRI adaptation paradigm to probe auditory and tactile frequency responses in the human brain. Adaptation describes the tendency for repeated sensory stimulation to alter the responses of neurons that are tuned for features of the sensory stimuli (Solomon and Kohn, 2014). fMRI adaptation paradigms leverage this process to infer population-level (i.e., voxel-wise) tuning properties based on BOLD signal changes (Grill-Spector et al., 2006). We scanned healthy human participants performing a covert frequency monitoring task on brief auditory, tactile, and audiotactile stimulus sequences that contained repeating or changing frequencies. In univariate analyses, we quantified BOLD signal changes related to unimodal (auditory-only or tactile-only) and crossmodal adaptation effects to identify brain regions exhibiting activation patterns consistent with population-level frequency tuning. We additionally performed multivariate pattern analysis to establish the representational spaces occupied by the unimodal and crossmodal events in different regions spanning the parietal and temporal lobes. We used representational similarity analysis (RSA) to determine the relationships between the neural subspaces of different brain regions. We then compared the cortical landscape defined by the event-related spatial activation patterns to the connectivity profile computed from temporal correlations between spontaneous signal fluctuations measured in the same participants in separate resting state scans. Collectively, our results indicate that frequency-selective auditory and tactile processing is broadly distributed over sensory systems which are organized according to the traditional modality-based scheme.

## Materials and Methods

### Participants

Twenty-four healthy adult participants were recruited from Baylor College of Medicine and the Houston metropolitan area. Participants indicated via self-report the absence of any current or past psychiatric or neurological conditions and were not taking any centrally acting medications. Participants reported normal tactile and auditory sensibilities. Testing procedures were performed in compliance with the policies and procedures of the Baylor College of Medicine Institutional Review Board. All participants provided informed written consent and were paid for their participation. A total for 4 participants were excluded from all analyses: 2 participants were excluded for below chance performance on the frequency monitoring task during the main adaptation scans (see below) and 2 participants were excluded for above chance performance on a control detection task (see below). All analyses thus included data from 20 participants (10m10f; mean age ± standard deviation: 23.2 ± 4.4 years). All participants were right handed according to the Edinburgh Handedness Inventory (mean score ± standard deviation: 79 ± 23.8) (Oldfield, 1971).

### Overview: Neuroimaging

Each participant underwent a functional localizer (Mapping) scan, 8 functional (Adaptation) scans involving continuous performance of a frequency monitoring task, 1 structural scan, and 2 resting state scans. Each participant also completed a control behavioral task in the scanner (Detection control).

### Image acquisition

All scans were conducted in the Core for Advanced MRI (CAMRI) at Baylor College of Medicine. MRI data were acquired on a Siemens MAGNETOM Trio 3 Tesla system using a Siemens 32-channel head coil. Structural images were acquired with a sagittal magnetization prepared rapid gradient echo (MP-RAGE) T1-weighted sequence (time echo (TE) = 3.02ms; time repetition (TR) = 2600ms; time to inversion = 900ms; flip angle = 8°; GRAPPA factor = 2; 176 slices with 1 × 1 × 1 mm voxels). Functional images for task runs were obtained using an axial echo-planar imaging (EPI) sequence (TE = 25ms; TR = 2750ms; flip angle = 80°; GRAPPA factor = 4; 56 slices; 2 mm^3^ voxels; 0 mm gap). A total of 151 volumes were acquired for the mapping scan, 121 volumes were acquired for each of the 8 adaptation scans, and 121 volumes were acquired for each of the 2 resting state scans.

### Auditory and tactile stimulation

Auditory stimuli were generated in Matlab (2011b; Mathworks) running on a Macbook and sounds were delivered via MRI-compatible in-ear headphones (Model S14, Sensimetrics) after amplification (PCA1, Pyle). Vibrotactile stimulation was delivered simultaneously to the distal finger pads on digits 2-5 on the right hand using a piezoelectric stimulator (CM3, <u>Cortical Metrics</u>). Because we tasked participants with monitoring the frequency of the auditory and tactile cues, all sounds and vibrations were matched for perceived intensity (Perez-Bellido et al., 2017). To account for individual differences in perceptual sensitivity and to ensure further that stimulus amplitude did not systematically vary with stimulus frequency, we introduced a random jitter (±5%) to the intensity-matched amplitude levels on every stimulus presentation.

### Modality-mapping localizer scan

Each subject underwent a functional scan designed to identify brain regions responsive to tactile or auditory stimulation (Mapping scan). Using a block design paradigm (Fig. 1), we presented unimodal blocks (16.5s) of tactile stimulation (25-400Hz), auditory pure-tone stimulation (40-700Hz), or auditory band-pass noise stimulation (center frequencies: 40-700Hz; bandwidths: 0.45*center frequency). Blocks were separated by 11s of no auditory or tactile stimulation. Each unimodal block consisted of 24 randomly ordered stimuli (duration: 437.5ms; inter-stimulus interval (ISI): 250ms). Each block type was repeated 5 times in pseudorandom order. Participants passively experienced the stimulation while maintaining fixation on a visual display.

**Figure 1.**
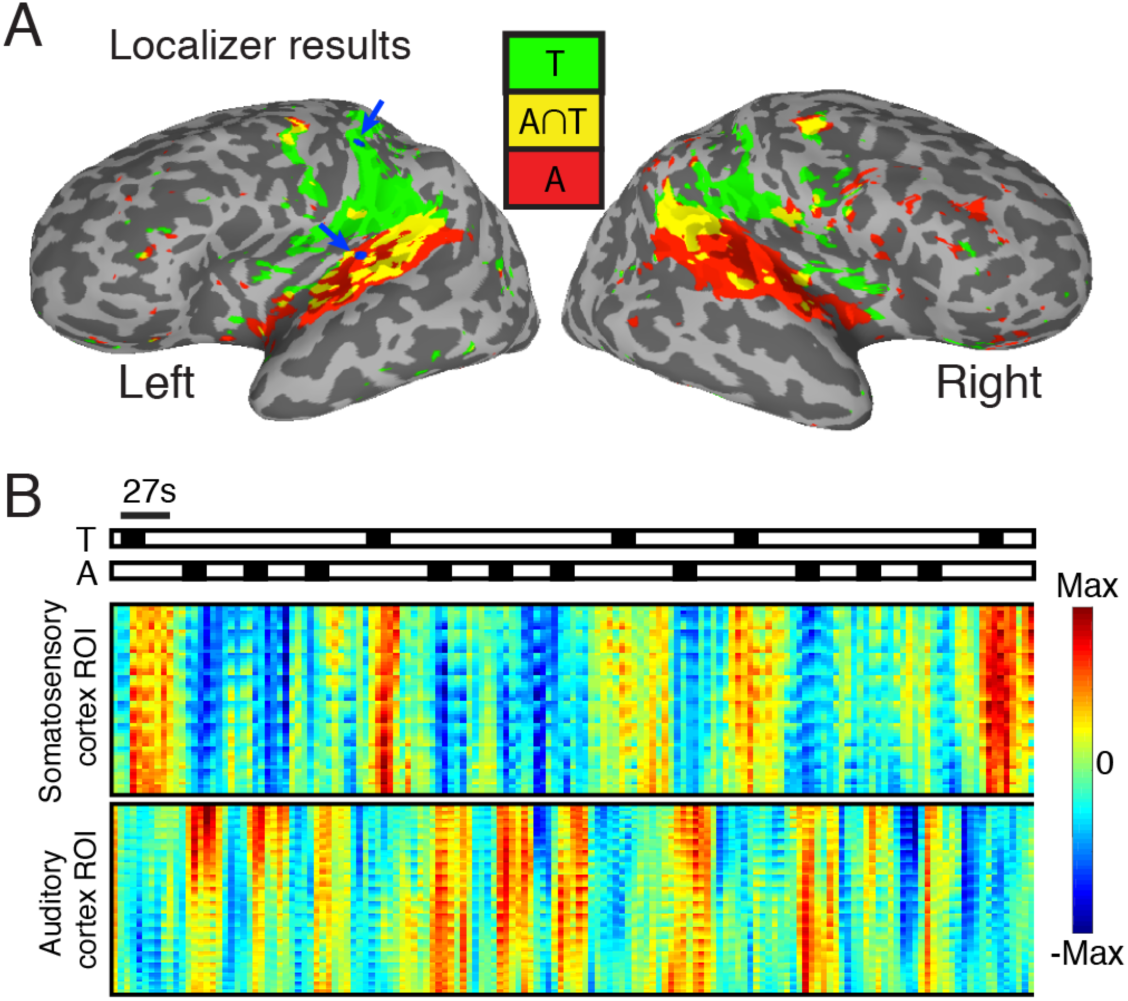
Localizer results. ***A***, Group results from block-design functional localizer scans (Materials and Methods) depicting regions that responded to only auditory stimulation (red; pure tones or band-pass noise) or only tactile stimulation (green) (*p* < 0.05, uncorrected). Yellow regions indicate the intersection of auditory and tactile activations. The blue arrows indicate parietal and temporal regions-of-interest (ROI, blue nodes) in the left hemisphere whose time series are depicted in *B*. The union of the auditory and tactile activations defines the functional mask used in the analysis of the event-related adaptation scans. ***B***, Stimulus time series for a localizer scan in an example participant depicting blocks during which tactile (T) or auditory (A) stimulation were delivered (Materials and Methods). Time series for somatosensory cortex ROI and auditory cortex ROI. Each row indicates normalized BOLD signals measured in a single surface node. Activity profiles in each ROI reveal clear response modulation associated with the sensory stimulation blocks.

### Frequency adaptation scans

Each subject underwent 8 functional scans testing frequency adaptation. Each scan comprised 48 adaptation events. Each event (Fig. 2A) consisted of 3 adapting stimuli followed by 1 probe stimulus. Stimuli (500-ms duration; 250-ms ISI) were either 100Hz or 300Hz. We tested 16 adaptation event types in a 2 × 2 × 2 × 2 factorial design (Fig. 2B) that varied the modality of the adapting stimuli (auditory or tactile), the modality of the probe stimulus (auditory or tactile), the frequency of the adaptors, and the frequency relationship between the adaptors and the probe (repeat or change). Thus, we tested within-modality adaptation sequences (auditory adaptors – auditory probes, AA; tactile adaptors – tactile probes, TT) and across-modality adaptation sequences (auditory adaptors – tactile probes, AT; tactile adaptors – auditory probes). Each event type was presented 6 times per scan (yielding a total of 48 repeats per event type over all scans). Half of the REPEAT trials contained an adapting frequency of 100Hz and the adapting frequency in the other half were 300Hz. CHANGE trials also contained an equal number of 100-and 300-Hz adaptors. This design ensured that the frequency and modality of the adapting stimuli did not predict the frequency or modality of the probe stimulus on any given event.

**Figure 2.**
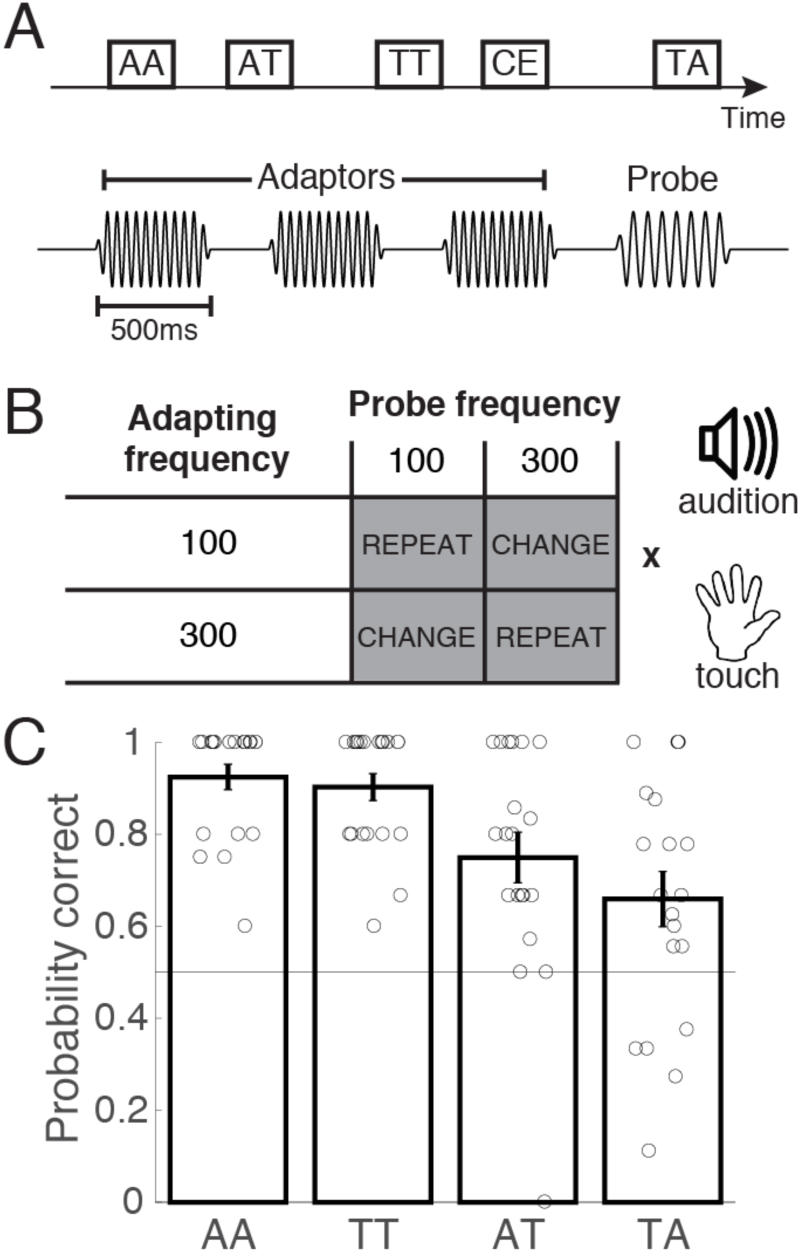
fMRI adaptation paradigm and behavioral results. ***A***, Event-related design comprising within-modality and across-modality adaptation events. Each 2.75-s event is a stimulus sequence comprising 3 adaptor stimuli that are matched in frequency and a probe stimulus whose frequency either repeats or changes with respect to the adapting frequency. The modality of the adaptors and probes can be matched (auditory-auditory, AA; tactile-tactile, TT) or different (auditory-tactile, AT; tactile-auditory, TA). Participants are tasked with monitoring the frequency of each stimulus and covertly determining if the frequency of the probe matches that of the adaptors. Participants report their decisions when prompted visually during catch events (CE). ***B***, Adapting and probe frequencies could be either 100Hz or 300Hz. The experiment comprised a full parametric design crossing adapting frequency, probe frequency relation to adapting frequency (REPEAT or CHANGE), and stimulus modality. ***C***, Average task performance for each event type. Circles indicate single participant performance levels (n = 20). Error bars indicates S.E.M.

Subjects fixated a central visual crosshair during each trial. To maintain participants’ attention on stimulus frequency throughout the scans, participants were asked to compare the frequencies of the adapting and probe stimuli on each trial and judge whether they were the same or different. However, participants only reported their judgments by button press when prompted on occasional catch events (3 per scan) with their left hand using an MRI-compatible button box (Current Designs). This design enabled us to monitor compliance on the frequency monitoring task without contaminating our estimates of sensory processing activity with motor response activity. Two participants failed to achieve above-chance performance over all of the conditions, so we excluded their data from additional analyses. The ordering and timing of events in each scan was optimized and pseudorandomized using Optseq2 (http://surfer.nmr.mgh.harvard.edu/optseq).

### Tactile detection control task

Because we were interested in comparing responses to auditory and tactile stimulation, it was imperative for us to verify that participants could not judge tactile stimulus frequency based on acoustic cues produced by the piezoelectric stimulator in the scanner. Accordingly, we tested each participant on a 2-interval forced-choice vibration detection task while the stimulator and scanner were operating but with the stimulator positioned next to the body without contacting the hand or any other body part. On each trial, a 500-ms tactile stimulus (25, 100, or 300Hz) was generated by the piezoelectric stimulator in one of two intervals (ITI = 1000ms). Participants were required to report the interval in which they detected the stimulus. Trial and response intervals were cued visually, and participants were instructed to guess if uncertain. We reasoned that participants would perform at chance level if they were unable to perceive the tactile stimulus without physical contact. Conversely, we reasoned that participants would perform above chance level if they exploited alternative cues (e.g., stimulator-generated sounds) to perform the tactile detection task. To replicate the acoustic conditions of the adaptation and mapping runs during the tactile detection task, we operated the scanner using the same EPI sequence. Two participants achieved above-chance performance on the detection task, so we excluded their data from additional analyses.

### Data analysis: Overview

We first defined brain regions that exhibited significant responses to auditory or tactile stimulation in analysis of the functional localizer scans (Fig. 1). Regions identified from the localizer data then constrained the data space in the analyses of the frequency adaptation scans, which included univariate and multivariate tests. All analyses were performed in surface space. As a first step in the analysis of the frequency adaptation scans, we conducted group-level statistical tests of parameters estimated in general linear models (GLM) fitted to the time series of each surface node. We supplemented this node-based analysis with region-of-interest (ROI) analyses that specifically focused on putative auditory and somatosensory regions in the temporal and parietal lobes defined using a combination of functional masks (i.e., the localizer results) and anatomical masks (see below). We conducted separate ROI-based analyses to evaluate adaptation effects, to compare representational geometries based on multivariate response patterns, and to relate brain regions based on their patterns of spontaneous BOLD signal fluctuations. Each analysis is described in detail in the following sections.

### fMRI data pre-processing

We used AFNI software (Cox, 1996) to perform data pre-processing and univariate analyses. Three-dimensional surface models were created with FreeSurfer (Fischl et al., 1999) (freesurfer-Darwin-lion-stable-pub-v5.3.0) and visualized in SUMA (Saad and Reynolds, 2012). Functional datasets were corrected for slice timing, motion corrected, and despiked. Volumes with head motion exceeding 0.3 mm/TR or a fraction of 0.05 outlier voxels (calculated using the 3dToutcount function) were excluded from analysis. Data from the localizer scan and the frequency adaptation scans that were included in whole-brain univariate analyses were normalized to the N27 atlas brain (Mazziotta et al., 2001) and spatially smoothed (4-mm FWHM 2D Gaussian kernel) in standard surface space. No warping or smoothing was applied to data from the frequency adaptation scans and the resting state scans that were included in the ROI-based analyses. All data were expressed in percent signal change with respect to the mean signal in each scan.

### fMRI data analyses: Localizer scans

Localizer scans were modeled using GLMs that included 3 regressors of interest corresponding to tactile stimulation, auditory pure-tone stimulation, and auditory band-pass noise stimulation convolved with gamma-variate functions. Head motion parameters and drift parameters (linear, quadratic, and cubic) were included as nuisance regressors. In group-level analyses, we identified the surface nodes that exhibited positive BOLD signal changes in each block type (Fig. 1). Because we only used these patterns to define an analysis mask, we used a liberal uncorrected threshold (*p* < 0.05). Activation maps for auditory and tactile stimulation blocks were combined to generate a single analysis mask.

### fMRI data analyses: Adaptation scans

We fitted node-wise multiple linear regression models in first-level analyses. The simplest GLM included 8 stimulus regressors (4 adaptor-probe modality conditions: AA, TT, AT, TA; 2 adaptation conditions: CHANGE, REPEAT) and catch events (CE) as separate regressors. This model (9 conditions of interest) assumed that there were no response differences related to adapting frequency. We also fitted a more complex GLM which allowed for effects of adapting frequency; this model included 16 adaptation conditions (4 modality conditions; 2 adaptation conditions; 2 adapting frequencies: 100Hz and 300Hz) along with CE for a total of 17 regressors of interest. Each regression model also included motion and drift parameters as nuisance regressors. Gamma-variate functions were used to model stimulus responses in the main analyses. Beta coefficients from the first-level analyses were submitted to group-level univariate analyses designed to identify nodes that exhibited significant response modulation related to the adaptation conditions. All group-level univariate test results were constrained by the mask defined by the localizer results and statistically thresholded at a false discovery rate (FDR) corrected *q* < 0.05. We used the ‘SurfClust’ AFNI function to summarize the clustered activation patterns identified as significant after FDR correction. In a separate validation procedure, we additionally analyzed the data using finite impulse response deconvolution in order to visualize response time courses for each stimulus condition (8 time points; 0-19.25s post-event onset).

### Within-modality adaptation events

We performed a single analysis on the within-modality adaptation events (AA and TT) by conducting a two-way repeated measures ANOVA (rmANOVA) with modality (auditory or tactile) and adaptation condition (CHANGE or REPEAT) as the factors. This analysis ignores potential modulating effects of adapting frequency. We evaluated main effects of modality and adaptation condition as well as the modality × adaptation interaction. For each node showing a significant main effect of adaptation condition, we defined an adaptation index (AI) as the response on CHANGE events minus the response on REPEAT events. Large absolute AI values indicate greater frequency-sensitivity and positive AI values reflect repetition suppression. For each participant, we sorted the data into 11 anatomically-defined ROIs (Destrieux et al., 2010) spanning the parietal and temporal lobes: central sulcus (cS), postcentral gyrus (pcG), postcentral sulcus (pcS), supramarginal gyrus (SMG), subcentral gyrus and sulcus (subcG/S), posterior Sylvian fissure (pSF), planum temporale (PT), transverse temporal sulcus (tTS), superior temporal gyrus (STG), lateral superior temporal gyrus (latSTG), and the inferior Insular cortex (infIns). To determine if AI values differed according to modality and ROI, we conducted a two-way rmANOVA with modality and ROI as factors.

### Across-modality adaptation events

We first tested for adaptation effects that were invariant to adapting frequency using the two-way rmANOVA with modality and adaptation condition as factors. Because no regions showed significant main or interaction effects of adaptation condition, we then tested for adaptation effects that depended on the adapting frequency. For the AT and TA conditions separately, we conducted two-way rmANOVA with adapting frequency (100Hz or 300Hz) and adaptation condition (REPEAT or CHANGE) as factors.

### Modeling across-modality adaptation responses

For activation clusters identified as significant for the adapting frequency × adaptation condition interaction, we evaluated a set of competing models that related β_Obs_ – the observed BOLD response profile over the frequency conditions (adaptor-probe: 100-100, 100-300, 300-100, 300-300) – to linear combinations of 4 potential predictor variables: adaptor frequency (f_a_), probe frequency (f_p_), response pattern to tactile-only events (β_TT_), and response pattern to auditory-only events (β_AA_). These terms enabled us to quantify how the adapting and probe frequencies contributed to the across-modality response pattern. We also reasoned that a component of a cluster’s across-modality responses may reflect its response to the within-modality events. Thus, we tested 4 hypotheses by fitting 4 alternative models:

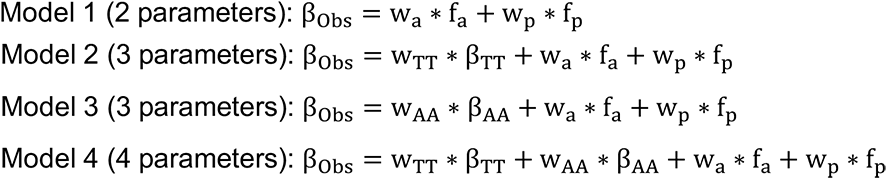

where, w_a_, w_p_, w_TT_, and w_AA_ are the weights to the model terms based on adapting frequency, probe frequency, the TT response pattern, and the AA response pattern. For each participant, the models were trained and tested on data over all of the significant clusters using a 10-fold cross-validation procedure: On each fold, the model parameters were estimated using 90% of the data and model performance was determined using the held-out data. The cross-validation procedure was repeated 1000 times to generate distributions of parameter estimates and goodness-of-fit values. We performed model comparison using Akaike information criteria (AIC) and Bayesian information criteria (BIC) to identify the preferred model.

### Representational similarity analysis (RSA)

We compared the spatial activation patterns associated with each event to define the representational space occupied by the adaptation stimulus sequences for each ROI. These analyses included any node identified as significant in the univariate analyses. Each participant’s scans were divided into 2 datasets on which separate GLMs were fitted to estimate 2 sets of multivariate patterns for each of the 17 event types (2 modalities × 2 adapting frequencies × 2 adaptation conditions + 1 catch event). We then computed correlations between the activation patterns associated with all possible pairs of events. We calculated a distance metric for each pair of events (1 minus the correlation) where distances ranged between 0 (identical pattern) and 2 (perfectly anti-correlated pattern). The full distance matrix (DM) thus defined the representational space for a given ROI. Note that separate analyses in which response pattern similarity was defined by Mahanalobis distance instead of correlation yielded similar overall results.

To quantify how the similarity of activation patterns in each ROI related to the modality of the stimuli comprising the events, we defined a tactile modality similarity index (MSI_T_) for each ROI using the 1^st^ order DMs:

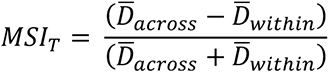

where 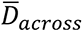 is the mean distance between the tactile-only events and the across-modality (AT and TA) events and 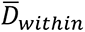 is the mean distance between just the tactile-only events. Larger MSI_T_ values indicate greater relative response pattern similarity among tactile-only events, which we expected for conventionally-defined somatosensory areas. We computed an analogous auditory modality similarity index (MSI_A_) for each ROI as well.

To establish the similarity of representational spaces across ROIs, we compared the 1^st^ order DMs established for the different ROIs and computed the correlations between the DMs of ROI pairs. We calculated a distance metric for each pair of ROIs (1 minus the correlation) to construct a 2^nd^ order DM where distances ranged from 0 (identical 1^st^ order DMs) and 2 (perfectly anti-correlated 1^st^ order DMs). The full 2^nd^ order DM thus defined the ROI landscape for each participant.

In group-level analyses, we averaged 2^nd^ order DMs over participants to establish a mean 2^nd^ order DM. We applied single-linkage hierarchical clustering on the group DM to quantitatively characterize the group-level ROI landscape. To better visualize the ROI landscape, we used classical multidimensional scaling (MDS) and projected the ROI distance matrix into a 2-dimensional subspace that maximally preserved the distances between ROI pairs. MDS was applied to each participant’s 2^nd^ order DM and the subspace representations were averaged after Procrustes alignment (Ejaz et al., 2015).

### ROI-based resting-state analysis

This analysis was aimed at characterizing the similarity of spontaneous BOLD signal fluctuations across the 11 parietal and temporal ROIs. After standard preprocessing, BOLD signal time series were estimated for each node by taking the residuals after regressing out head motion and physiological noise signals (Jo et al., 2010; Barnes et al., 2014). Time series were extracted for all of the nodes included in the RSA. For each participant, a mean time series was calculated for each ROI separately. The correlation between the mean time series of pairs of ROIs was calculated and the correlations were converted into distances (1 minus correlation) ranging from 0 (perfectly correlated time series) and 2 (perfectly anti-correlated time series). This distance matrix thus provided an ROI landscape defined according to temporal correlations in participants’ resting state data. Distance matrices were averaged across participants for a group DM which was directly compared to the group-averaged 2^nd^ order DM calculated from the RSA. We computed the correlation between the group DMs to determine the similarity of the ROI landscapes determined according to the spatial activation profiles in the task-based data and the temporal correlations present in the resting state data. We tested the significance of this correlation using a randomization test which compared the observed correlation against a null distribution of correlations expected by chance. The null distribution was computed by repeating the correlation calculation 1000 times with the node-wise data randomly assigned over the ROIs.

## Results

### Behavioral results: Performance levels are consistent with participants attending to stimulus frequency

Participants performed a frequency monitoring task as they underwent fMRI. Each event comprised a series of 4 brief stimuli delivered to the participant’s right hand and participants covertly judged whether the frequency of the probe stimulus matched the frequency of the preceding three adaptor stimuli. To confirm that participants performed this covert frequency monitoring task, participants were required to report their frequency judgment by button press when visually cued randomly throughout each scan. Participants generally maintained high performance on this task (Fig. 2C). Although performance varied significantly according to the modality conditions (F = 12.4, P = 2.3e-6), performance levels on average exceeded chance (0.5) on both the within-modality trials (AA: 0.93 ± 0.03, T = 15.3, P = 3.8e-12; TT: 0.90 ± 0.03, T = 13.8, P = 2.3e-11) and across-modality trials with auditory adaptation (AT: 0.75 ± 0.06, T = 4.5, P = 2.3e-4). Performance on across-modality trials with tactile adaptation was more variable across subjects and the group average was marginally greater than chance (TA: 0.66 ± 0.06, T = 2.7, P = 0.016). These results indicate that most participants were compliant in attending to stimulus frequency during the unimodal and crossmodal events.

### Within-modality repetition suppression reveals frequency-selective auditory and tactile processing in overlapping regions

We identified a number of regions that showed a main effect of stimulus modality (Table and the modality preference map closely resembled the localizer results. To identify brain regions whose response patterns were consistent with population-level (voxel-wise) neural selectivity for auditory or tactile frequency, we contrasted responses measured when the probe frequency differed from the adapting frequency (CHANGE events) against responses measured when the probe frequency matched the adapting frequency (REPEAT events). This contrast revealed significant activation clusters (Fig. 3A) that were largely confined to the parietal and temporal lobes in the left hemisphere (Table 1). Notably, these clusters all exhibited greater responses to CHANGE events compared to REPEAT events, revealing a pattern consistent with repetition suppression. This pattern is evident in the GLM coefficients (Fig. 3B) and estimated temporal response profiles (Fig. 3C) for auditory-only and tactile-only events in an example ROI, the posterior Sylvian fissure. In fact, greater responses to CHANGE events compared to REPEAT events were observed in nearly all of the defined temporal and parietal ROIs with the auditory-only events (Fig. 4A) and tactile-only events (Fig. 4B). Although auditory-only and tactile-only events were both associated with distributed responses over putative auditory and somatosensory regions, auditory responses were relatively greater in temporal regions while tactile responses were relatively greater in parietal regions. Importantly, no regions showed significant modality × adaptation interaction effects, indicating that adaptation response patterns tended to be consistent for the AA and TT events.

**Figure 3.**
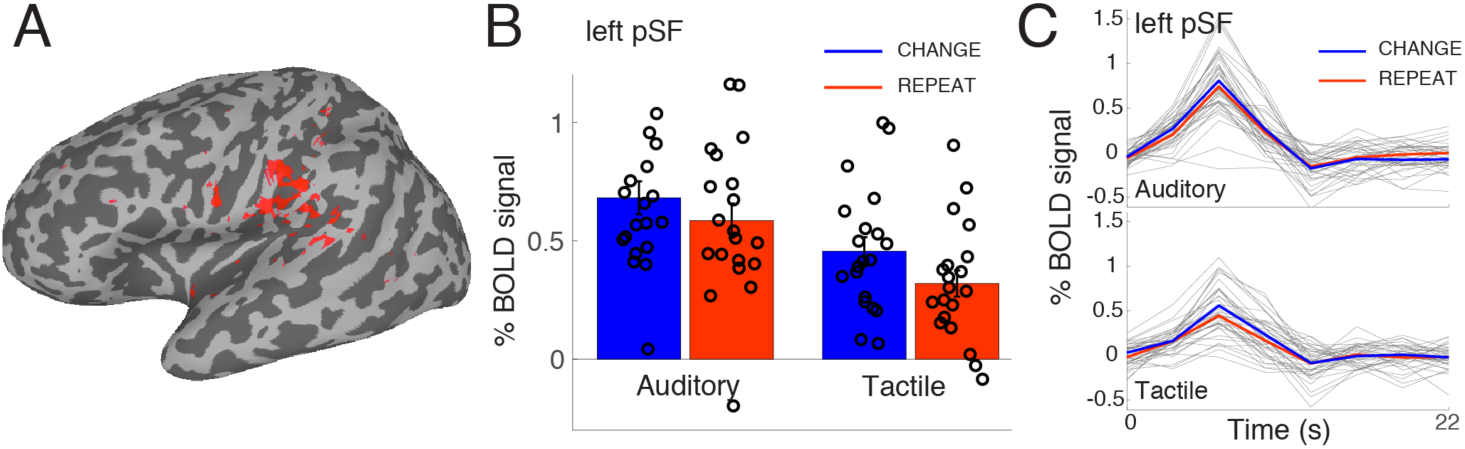
Within-modality adaptation results. (n = 20). ***A***, Red nodes indicate regions that exhibited significant response differences between CHANGE and REPEAT events regardless of modality (FDR corrected, *q* < 0.05; constrained by functional localizer mask). ***B***, Group-averaged response magnitudes estimated for auditory and tactile within-modality adaptation events for the left posterior Sylvian fissure region-of-interest (pSF). This ROI exhibited responses to both auditory-only and tactile-only stimulus sequences. CHANGE events (blue bars) were associated with greater responses compared to REPEAT events (red bars) for auditory and tactile comparisons. Error bars indicate S.E.M. ***C***, Group-averaged temporal response profiles estimated for auditory and tactile within-modality adaptation events for left pSF. Responses to CHANGE events (blue traces) tended to larger than responses to REPEAT events (red traces). Thin traces indicate individual subject data.

**Figure 4.**
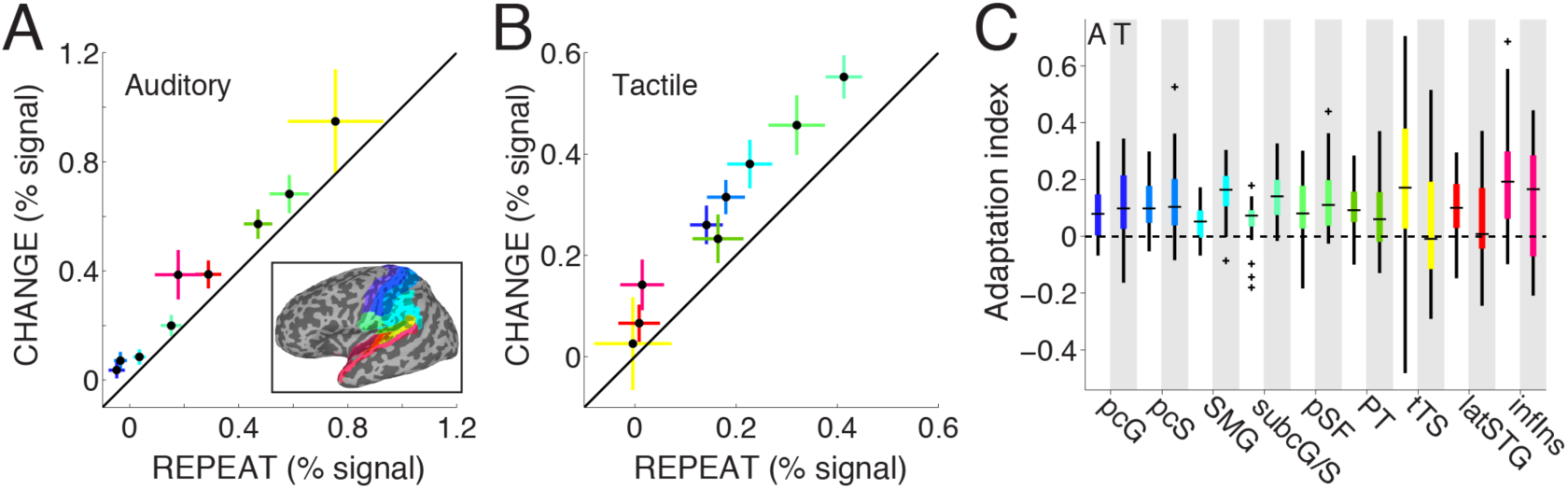
ROI-based within-modality adaptation results. (n = 20) ***A***, Average responses to auditory-only REPEAT and CHANGE events. Each marker indicates mean response over nodes in different parietal and temporal ROIs (inset) exhibiting a significant main effect of adaptation condition (Fig. 3A). Error bars indicate S.E.M. ***B***, Average responses to tactile-only REPEAT and CHANGE events. Conventions as in *A*. ***C***, Box and whisker plot depicts adaptation index (Materials and Methods) for auditory-only (white columns) and tactile-only events (gray columns) for different ROIs. Horizontal black lines indicate medians. Box edges indicate interquartile range (IQR). Whiskers extend to most extreme data points within 1.5x IQR and individual points indicate data outside of 1.5x IQR. pcG, postcentral gyrus; pcS, postcentral sulcus; SMG, supramarginal gyrus; subcG/S, subcentral gyrus and sulcus; pSF, posterior silvian fissure; PT, planum temporale; tTS, tranverse temporal sulcus; latSTG, lateral superior temporal gyrus; infIns, inferior insula

**Table 1.** Univariate analysis: Regions that exhibited significant effects on the within-modality adaptation events

| # Nodes | Total area (mm<sup>2</sup>) | Maximum (F) value | Peak node | Talarach Peak coordinates (LPI) |  |  | Destrieux Atlas labels |
| --- | --- | --- | --- | --- | --- | --- | --- |
|  |  |  |  | x | y | z |  |
| Modality |  |  |  |  |  |  |  |
| Left Hemisphere |  |  |  |  |  |  |  |
| 6310 | 1368.06 | 4.772 | 146030 | -48 | -31 | 12 | Planum temporale |
| 5870 | 1344.73 | 4.761 | 149591 | -54 | -24 | 19 | Inferior parietal gyrus, supramarginal |
| 630 | 153.52 | 4.772 | 131648 | -43 | -5 | 13 | Circular sulcus of the insula, inferior |
| 289 | 100.24 | 3.874 | 85638 | -42 | -1 | 44 | Precentral gyrus |
| 362 | 93.95 | 3.948 | 96105 | -48 | -4 | 16 | Precentral sulcus, inferior |
| 125 | 60.58 | 3.765 | 13655 | 35 | 0 | -12 | Circular sulcus of the insula, inferior |
| 116 | 54.9 | 2.843 | 15563 | -36 | -7 | -9 | Circular sulcus of the insula, inferior |
| 88 | 37.15 | 4.257 | 40433 | -11 | -27 | -2 | UNDEFINED |
| 107 | 33.40 | 3.961 | 15009 | -32 | -3 | 12 | Circular sulcus of the insula, superior |
| 112 | 25.55 | 3.14 | 96645 | -56 | -7 | 11 | Subcentral sulcus/ gyrus |
| 53 | 18.43 | 4.4 | 14576 | -36 | -1 | 6 | Circular sulcus of the insula, superior |
| 54 | 17.98 | 2.649 | 140077 | -47 | -47 | 0 | Superior temporal sulcus |
| 63 | 15.89 | 2.702 | 72680 | -13 | -9 | 60 | Superior frontal gyrus |
| 40 | 15.60 | 2.219 | 10898 | -30 | 8 | -7 | Short insular gyrus |
| 15 | 12.6 | 2.514 | 3762 | -22 | 23 | -9 | H-shaped orbital sulcus |
| 67 | 12.06 | 3.381 | 125334 | -33 | -37 | 32 | Postcentral sulcus |
| 17 | 10.65 | 2.691 | 177367 | -51 | -62 | 4 | Middle temporal gyrus |
| 53 | 10.46 | 3.508 | 127412 | -40 | -28 | 37 | Postcentral sulcus |
| 34 | 10.06 | 2.494 | 83802 | -41 | -7 | 48 | Precentral gyrus |
| Adaptation condition |  |  |  |  |  |  |  |
| Left Hemisphere |  |  |  |  |  |  |  |
|  |  |  |  | x | y | z |  |
| 1108 | 254.53 | 2.543 | 130263 | -50 | -20 | 16 | Inferior parietal gyrus, supramarginal |
| 244 | 41.27 | 2.465 | 133511 | -50 | -26 | 17 | Inferior parietal gyrus, supramarginal |
| 140 | 33.01 | 2.459 | 145311 | -54 | -32 | 14 | planum temporale |
| 114 | 29.2 | 2.465 | 95565 | -48 | -7 | 16 | Precentral gyrus |
| 26 | 13.1 | 2.392 | 13775 | -34 | 0 | 8 | long insular gyrus and central insular sulcus |
| 96 | 12.54 | 2.465 | 146956 | -46 | -42 | 18 | Posterior segment of the lateral fissure |
| 84 | 10.97 | 2.282 | 149265 | -59 | -29 | 19 | inferior parietal gyrus, supramarginal |
| 32 | 10.93 | 2.15 | 95687 | -58 | -10 | 14 | Subcentral sulcus/ gyrus |

Significant clusters identified in group-level analyses ( $q < 0.05$ FDR corrected). Data are shown for subset of clusters containing 30 or more nodes. A total of 32 clusters (10 nodes minimum) exhibited a significant main effect of modality (AUDITORY vs TACTILE). A total of 23 clusters (10 nodes minimum) exhibited a significant main effect of adaptation condition (CHANGE vs REPEAT).

We compared the magnitude of response differences between CHANGE and REPEAT unisensory events by calculating an adaptation index for each sensory modality over the 9 ROIs comprising significant clusters (Fig. 4C). AI values differed significantly according to brain region (2-way rmANOVA; ROI main effect: F = 2.12, P = 0.04; ROI × modality interaction effect: F = 3.63, P = 0.0007), but the main effect of modality failed to achieve statistical significance (F = 0, P = 0.94). Because of the significant interaction effect, we compared adaptation indices for the auditory and tactile events in each ROI and found significant differences only in the supramarginal gyrus (T = 4.09, P = 2.2e-4) and the subcentral gyrus/sulcus (T = 3.14, P = 0.003) with larger adaptation effects observed for the tactile events in both ROIs. We additionally determined if there was a general relationship between auditory and tactile adaptation strength across temporal and parietal regions and found that adaptation indices for the two sensory modalities, averaged over participants, tended to be negatively correlated (mean ρ = −0.084 ± 0.026) (one-sample t-test versus 0; T = 3.212, P = 0.0048). When performed on each ROI separately, this analysis revealed only significant correlations in the supramarginal gyrus (mean ρ = −0.122 ± 0.038; T = 3.208, P = 0.0046). These results indicate that the neural populations most sensitive to auditory and tactile temporal modulation effects do not necessarily reside in the same voxels.

### Across-modality adaptation events reveal frequency-selective temporal modulation effects in parietal cortex

To identify brain regions whose response patterns were consistent with population-level crossmodal frequency adaptation, we contrasted CHANGE and REPEAT responses in analyses that combined the AT and TA events (i.e., analysis independent of adapting modality) and analyses performed on the AT and TA events separately. These contrasts failed to reveal any significant clusters. We then tested the possibility that across-modality interaction effects varied according to the adaptor and probe frequencies. Accordingly, for the AT and TA events separately, we performed contrasts that involved interactions between the adaptation condition (CHANGE or REPEAT) and adapting frequency (100Hz or 300Hz) factors. Although no clusters displayed significant effects for this interaction in analysis of AT events, we identified 15 significant clusters in anterior parietal cortex and lateral parietal cortex with TA events (Fig. 5A). The response profile in these clusters revealed a clear interaction pattern of distinct repetition effects which depended on adapting frequency (Fig. 5B). We observed repetition facilitation with 100-Hz adaptors as REPEAT events were associated with greater responses compared to CHANGE events. In contrast, we observed repetition suppression with 300-Hz adaptors as responses to REPEAT events were reduced relative to CHANGE events.

**Figure 5.**
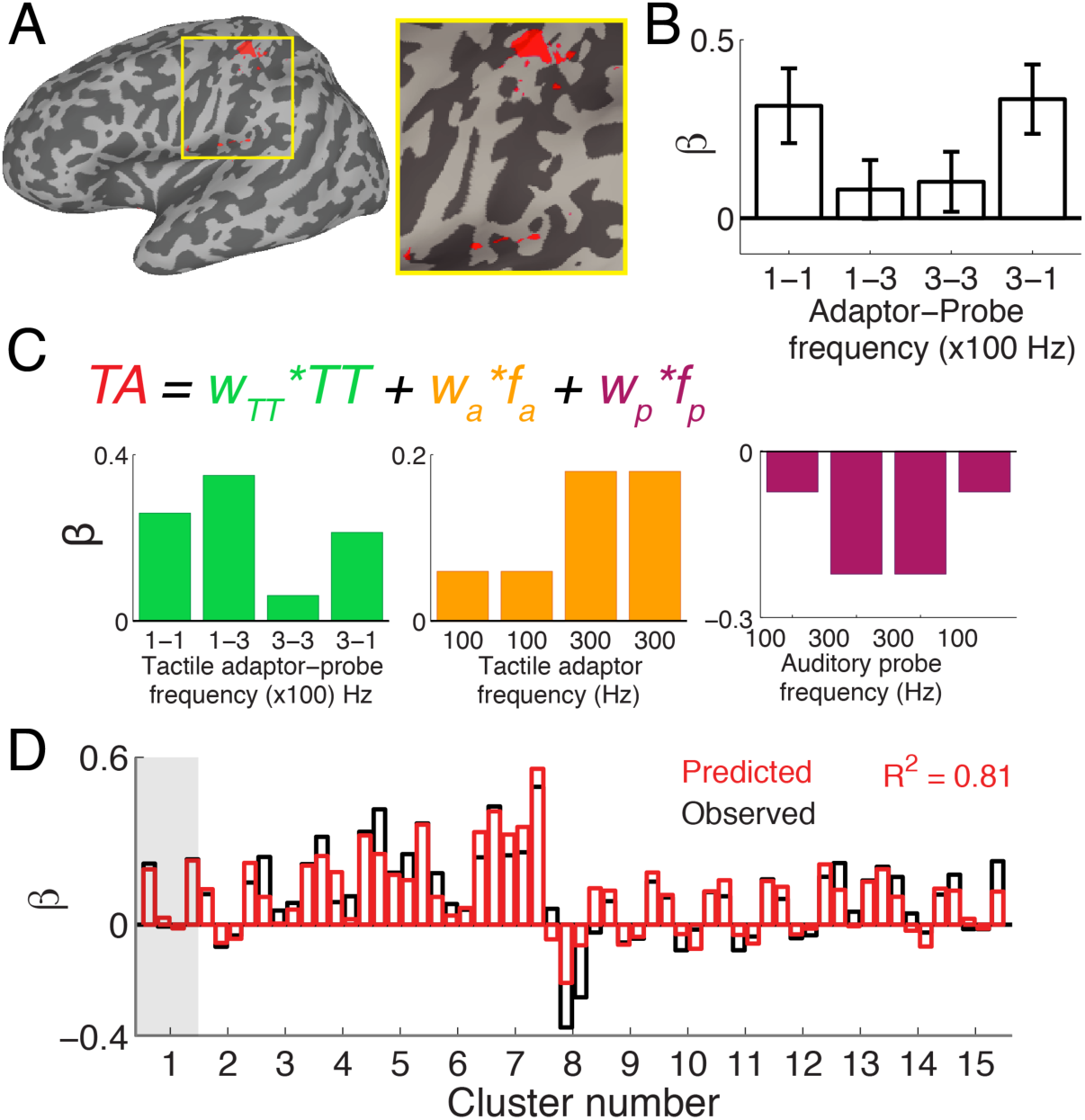
Results with tactile adaptation and auditory probes. (n = 20) ***A***, Crossmodal adaptation sequences comprising tactile adaptors and auditory probes (TA events) were associated with response modulation patterns in anterior and lateral parietal cortex (inset) that were characterized by significant interactions between adapting frequency (100Hz vs 300Hz) and adaptation condition (CHANGE vs REPEAT). ***B***, Average activation profile for example cluster. With 100-Hz adaptors, larger responses are observed on REPEAT events (i.e., with the 100-Hz probes compared to the 300-Hz probes). In contrast, with 300-Hz adaptors, larger responses are observed on CHANGE events (i.e., with the 100-Hz probes compared to the 300-Hz probes). Error bars indicate S.E.M. ***C***, TA response profiles in each cluster can be explained as a linear combination of the cluster’s TT response profile (green bars), frequency-specific response modulation attributed to the tactile adaptor stimuli (orange bars), and frequency-specific response modulation attributed to the auditory probe stimuli (purple bars) (Materials and Methods). The weighted sum of the depicted response components accounts for the activity pattern shown in *B*. ***D***, Observed and model predicted response patterns over 15 clusters exhibiting significant TA interaction patterns. Each cluster’s profile includes 4 responses (gray bar indicates profile for Cluster 1). Cross-validated models fit to response patterns in 90% of the data accounted for over 80% of the response variance in the held-out data (Materials and Methods).

To better understand the TA interaction pattern, we fit a set of simple linear models to the TA responses. Model comparison using both AIC and BIC identified the same preferred model (Fig. 5C) which comprised terms related to frequency-specific effects of the tactile adaptors and auditory probes as well as the TT response patterns. Figure 5D shows that the preferred model, fitted to responses over all 15 clusters exhibiting significant TA interactions in a cross-validated manner (Materials and Methods), explained greater than 80% of the response variance across clusters. Inspection of the model weights, capturing adaptor (0.000619 ± 0.000048) and probe (−0.00076 ± 0.0000382) frequency effects as well as the impact of the TT response profiles (0.76 ± 0.04), reveals that 1) the tactile adaptors contributed positively to cluster responses with larger modulations associated with the 300-Hz adaptors, 2) the auditory probes contributed negatively to cluster responses with larger modulations associated with the 300-Hz probes, and 3) the scaled TT response profile comprised a portion of the TA response profile. These results, along with the fact that no constraints were placed on which clusters contributed model training data and testing data, reveal that TA response patterns were highly consistent across the distributed clusters spanning anterior and lateral parietal cortex. In sum, the model results reveal that TA response patterns in parietal regions can be understand as a combination of repetition suppression (represented by the TT pattern), frequency-specific response enhancement from the tactile stimulus component, and frequency-specific auditory response inhibition from the auditory stimulus component.

### Spatial activation patterns are consistent with modality-based cortical organization

We performed multivariate analyses to further explore how unimodal and crossmodal adaptation sequences were represented across temporal and parietal regions (Material and Methods). In each ROI separately, we determined the pairwise similarity of spatial activation patterns associated with each event type in a split-half cross-validated manner (Fig. 6A). Figure 6B shows a 1^st^ order distance matrix calculated for the superior temporal gyrus in a representative participant. The elements of this DM describe the response pattern similarity between event pairs and the collective DM can be considered as reflecting the representational space occupied by the unimodal, crossmodal, and catch-trial events in STG neural space. The STG DM clearly shows that the spatial activation patterns associated with events comprising auditory stimuli (i.e., the AA, AT, TA events) tended to be highly similar to each other and dissimilar from TT events. Patterns associated with TT events also exhibited a degree of similarity with each other. While the representational space of STG is unsurprising given the known role of STG in auditory processing, the modest similarity between TT events reinforces the idea that temporal regions also respond systematically to tactile inputs alone.

**Figure 6.**
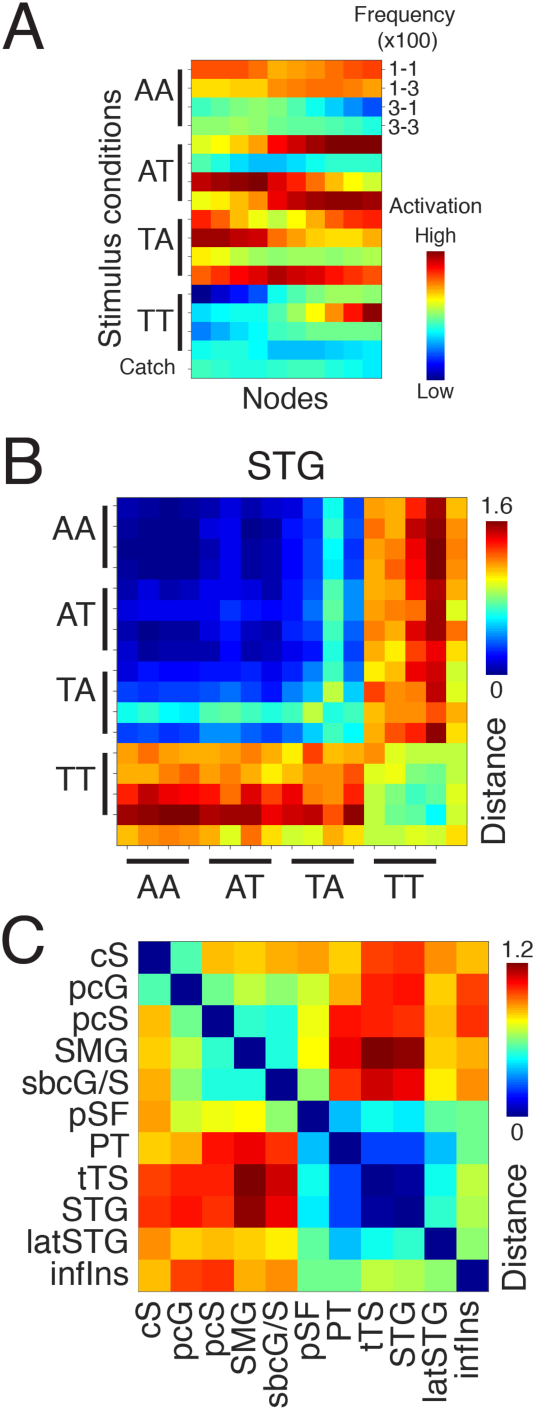
Representational structure of unimodal and multimodal stimulus sequences. ***A***, Activity matrix depicts the spatial activation patterns associated with each event (rows) over the nodes comprising a ROI (columns). Rows are additionally ordered according to frequency conditions (adaptor-probe frequencies) in each modality set (AA, AT, TA, and TT modality conditions). ***B***, First-order distance matrix (DM) indicating the pair-wise similarity between activation patterns associated with different event types in the superior temporal gyrus ROI. Lower distance values indicate more similar activation patterns. Similarity was quantified in a cross-validated manner with a single participant’s data acquired in different runs (Materials and Methods). Due to run-specific noise, identical events over different runs were associated with slightly different activation patterns which resulted in non-zero distances along the main diagonal. ***C***, Second-order DM in example participant indicating the similarity of first-order DMs between different ROI pairs. Lower distance values indicate that greater similarities in the representational spaces of two ROIs.

To establish the relationship between representational spaces in different temporal and parietal regions, we generated 1^st^ order DMs for each ROI and subsequently quantified the similarity of these DMs between ROI pairs. A 2^nd^ order DM for the representative participant is shown in Fig. 6C, with each element describing the similarity of the representational spaces of ROI pairs. The example DM reveals systematic relationships between representational spaces over the ROIs. First, there were greater similarities among the regions contained within the parietal lobe and the regions within the temporal lobe. Second, the representational spaces of parietal regions tended to be dissimilar from those of temporal regions. This systematic pattern was also obvious in the average 2^nd^ order DM over participants (Fig. 7A) with hierarchical clustering revealing 2 ROI groupings that were clearly organized according to regional affiliation to parietal or temporal cortex. To better visualize the relationship between ROIs, we projected and aligned each participants’ 2^nd^ order DM in a two-dimensional “brain region” subspace (Material and Methods). The configuration of the brain regions in this subspace (Fig. 7B) clearly reveals structured relationships over the ROIs with a parietal “stream” – spanning anterior parietal cortex (central sulcus, postcentral gyrus, postcentral sulcus), lateral parietal cortex (subcentral gyrus and sulcus), and posterior parietal cortex (supramarginal gyrus) – and a separate temporal “stream” – spanning STG, tranverse temporal sulcus, lateral STG, planum temporale, and posterior Silvian fissure. Notably, the MDS pattern also indicates that the inferior insular cortex contains a representational space that is distinct from the other ROIs. To establish a better intuition for the range of representational spaces observed over the ROIs, we highlighted the average 1^st^ order DMs for the postcentral gyrus and the superior temporal gyrus (Fig. 7C). We also highlighted the average 1^st^ order DM for the planum temporal ROI, which occupied a location between the parietal and temporal groups in the brain region subspace. As expected, the representational spaces in the parietal and temporal regions reflected a clear modality-dependence: the postcentral gyrus DM showed greater similarities among events comprising tactile cues (TT, TA, AT) while the STG DM showed greater similarities among events comprising auditory cues, as seen in the example participant (Fig. 6B). In contrast, because sensory events were generally associated with more similar activity patterns in planum temporale, the DM computed for PT was less characterized by modality-dependence. This pattern implies that regions like the PT and pSF, which are traditionally considered to be higher-order sensory association areas, differentiate between the auditory and tactile stimulus components less in their responses to the unimodal and crossmodal events than regions like the pcG and STG. To quantify these ROI differences, for each ROI we defined separate indices that expressed the modality-dependence of response pattern similarity and how response magnitudes varied according to modality (Materials and Methods). Both metrics (Fig. 8) varied significantly over ROIs (MSI: main effect of ROI: F = 6.5, P = 1.3e-8, main effect of modality: F = 21.2, P = 0.0002, ROI × modality interaction: F = 28.2, P = 2.2e-16; Response magnitude: main effect of ROI: F = 25.8, P = 2.2e-16, main effect of modality: F = 134.5, P = 4.6e-10, ROI × modality interaction: F = 85.1, P = 2.2e-16). These indices reveal cortical landscapes that are consistent with the conventional view of sensory systems: early sensory areas are more dedicated to a single modality and higher-order association areas are responsive to multiple modalities. Thus, the multivariate results are consistent with the traditional notion of modality-based sensory systems.

**Figure 7.**
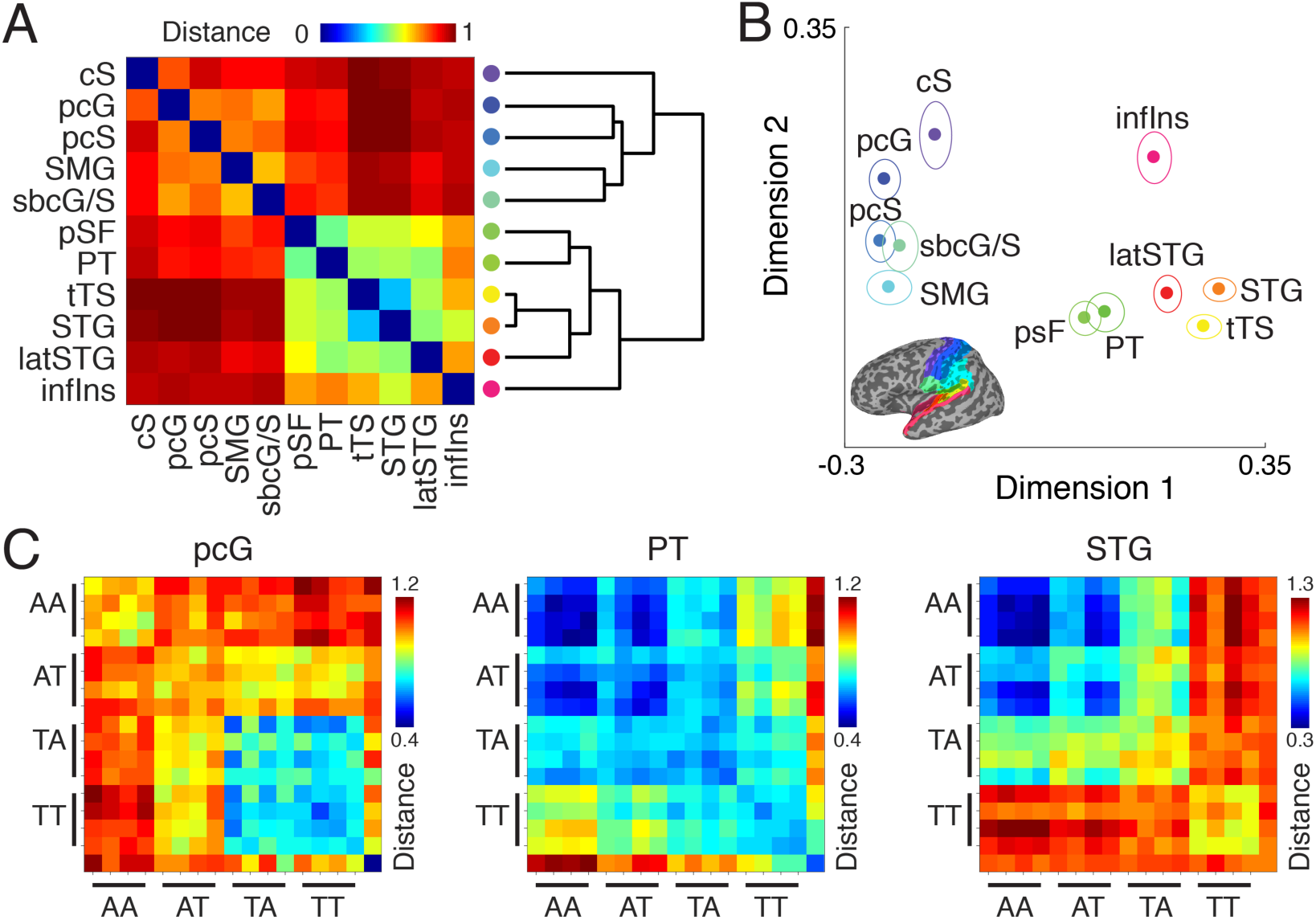
Representational spaces over parietal and temporal cortex. (n = 20) ***A***, Mean second-order distance matrix (DM) indicating similarity of representational spaces in different ROIs. Lower distance indicates greater similarity. Dendrogram depicts the clustering of parietal and temporal regions based on similarity of representational spaces. ***B***, Multidimensional scaling of the ROI distances in two-dimensional space. Ellipses show S.E.M. after Procrustes alignment across participants. ***C***, Mean representational spaces (first-order DM) for example regions occupying different portions of the ROI landscape shown in *B*. The DMs for postcentral gyrus (pcG) and superior temporal gyrus (STG) contain patterns clearly reflecting strong modality preferences for tactile and auditory stimulus components, respectively. The DM for planum temporale (PT) indicates spatial activation patterns that are more modality-invariant.

**Figure 8.**
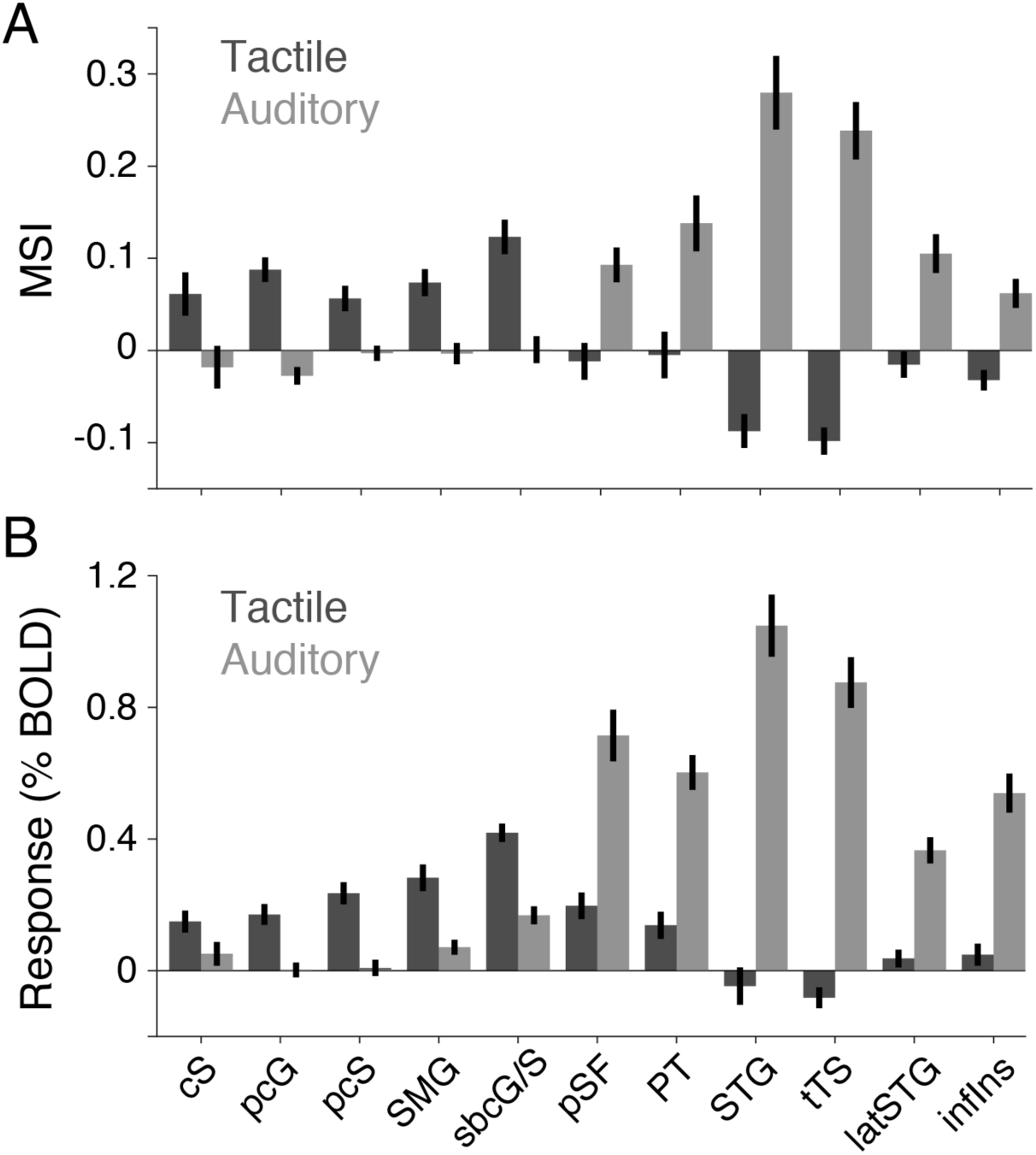
Region-of-interest landscapes. (n = 20) ***A***, Modality similarity index (MSI) defined according to spatial activation patterns in each ROI (Material and Methods). Parietal ROIs tended to have greater tactile MSI values (dark gray bars) while temporal ROIs tended to have greater auditory MSI values (light gray bars). Error bars indicate S.E.M. ***B***, Average response magnitude over tactile-only (TT) events (dark gray bars) and auditory-only (AA) events (light gray bars). Error bars indicate S.E.M.

### Resting state functional connectivity patterns are also consistent with modality-based cortical organization

We next determined whether the structured relationships between the parietal and temporal regions, established from the similarity of representational spaces in each ROI, were intrinsic characteristics of our participants’ brains or merely a reflection of task-evoked responses given our particular paradigm and stimulus conditions. Accordingly, we performed an analysis on each participant’s resting state fMRI data that was analogous to the RSA performed on their task-based fMRI data. For each participant, we computed the average BOLD signal time series in each of the 11 temporal and parietal ROIs (Fig. 9A). We then calculated the correlation between the time series in each ROI pair to generate a “connectivity” matrix expressing the degree of similarity in the temporal variations of spontaneous signal fluctuations across the ROIs. The average connectivity matrix over participants was significantly correlated with the group-averaged 2^nd^ order DM defined in the RSA (Fig. 9B; ρ = 0.28, P < 0.001; mean null ρ = 0.16 ± 0.03). In fact, direct comparisons between the connectivity matrix and 2^nd^ order DMs within participants yielded correlations ranging from 0.84 to 0.95 with a significant mean correlation of 0.92 ± 0.03. Indeed, projecting and aligning the brain regions across participants in a two-dimensional subspace (Materials and Methods) resulted in a ROI landscape (Fig. 9C) that closely resembled the landscape generated in the multivariate pattern analysis. The close correspondence between the RSA results (based on spatial variations) and the resting state connectivity analysis results (based on temporal variations) implies that the modality-based organization of regions spanning parietal and temporal cortex is an intrinsic characteristic rather than a structured pattern that emerged as a consequence of our experimental manipulations.

**Figure 9.**
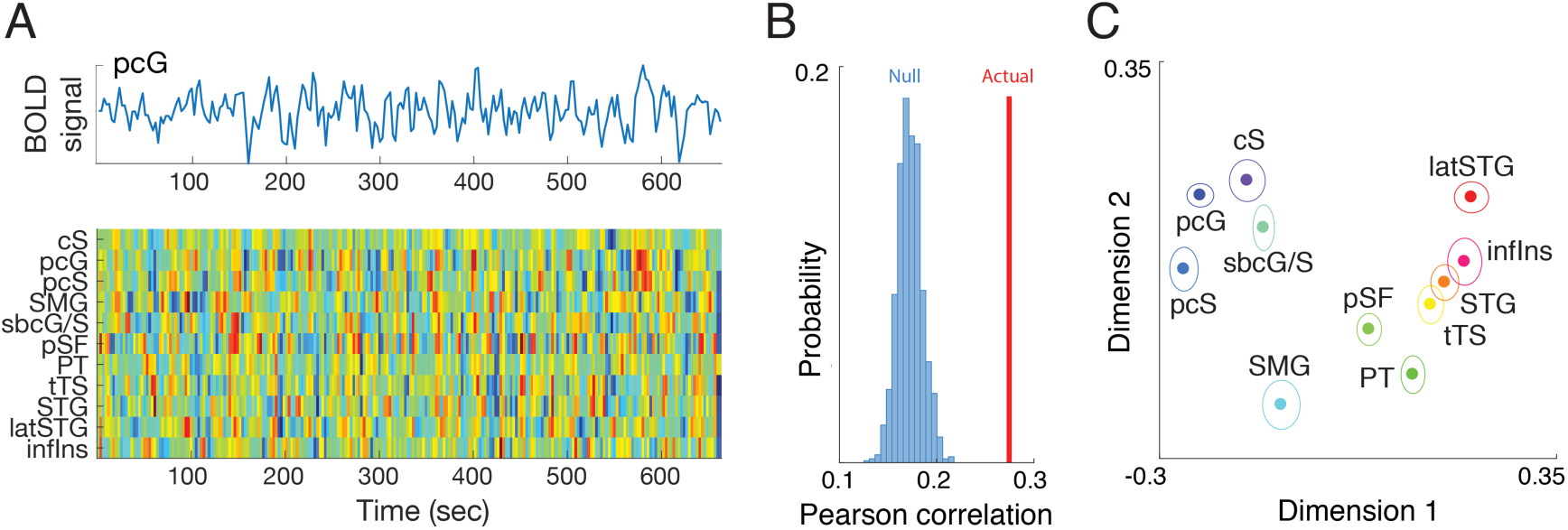
Region-of-interest relationships based on spontaneous BOLD signal fluctuations. (n = 20) ***A***, For each subject, mean time series were generated for each ROI from resting state scans. The time series for postcentral gyrus (top) is shown along with the time series for 10 other ROIs (below, rows). ***B***, Correlations between the time series of ROI pairs defined a second-order “connectivity” matrix that summarized the network architecture in the resting state data of each participant. This architecture was significantly correlated with the second-order distance matrix computed from representational similarity analyses (Fig. 7A): The actual correlation (red) between ROI landscapes generated from task-based data and resting state data far exceeded the correlations in the null distribution (blue). ***C***, Multidimensional scaling of the ROI distances (based on resting state correlations) in two-dimensional space. Ellipses show S.E.M. after Procrustes alignment across participants. Although the resting state analyses were performed independently of the task-based multivariate analyses, these yielded highly similar ROI landscapes.

## Discussion

We used an fMRI adaptation paradigm to identify brain regions exhibiting selectivity for auditory and tactile temporal frequency. With both unimodal tactile and auditory stimulus sequences, we observed repetition suppression in temporal regions, which we assumed would exhibit frequency selectivity *a priori*, as well as a number of parietal regions traditionally associated with somatosensory functions. To establish evidence for shared neural representations and multimodal frequency tuning, we tested crossmodal adaptation sequences and only observed significant frequency-dependent activity in parietal cortex for sequences comprising tactile adaptors followed by an auditory probe. Collectively, our adaptation results imply that both tactile and auditory frequency information modulate parietal and temporal cortex activity. We additionally analyzed the spatial activation patterns associated with the adaptation sequences in the parietal and temporal regions and found relationships between the ROI representational spaces that clearly reflected modality-based organization. The RSA results revealed a cortical landscape characterized by a set of modality-preferring regions linked through regions that were more modality-invariant. This landscape was also observed in the modality-sorted response amplitude profiles across the ROIs as well as the spatiotemporal patterns in spontaneous signal fluctuations measured in these areas at rest. Thus, although we found evidence for frequency-specific auditory and tactile processing distributed over parietal and temporal cortex, our data also support a traditional modality-based sensory cortex model.

The within-modality adaptation events were associated with BOLD signal changes that were consistent with repetition suppression effects (Grill-Spector et al., 2006; Krekelberg et al., 2006; Barron et al., 2016): Responses to stimulus sequences comprising repeats of a single frequency were significantly reduced relative to responses to sequences in which the probe frequency changed from the adaptor frequency. Many of the regions exhibiting frequency-based repetition suppression have also been shown to exhibit BOLD adaptation effects in other somatosensory (Hegner et al., 2007; Tame et al., 2012) and auditory (Millin et al., 2018) contexts. Our BOLD adaptation effects could relate to perceptual adaptation effects reported for frequency perception by somatosensation (Hollins et al., 1990; Goble and Hollins, 1993, 1994; Tommerdahl et al., 2005) and audition (Zwicker, 1964; Parra and Pearlmutter, 2007; Alais et al., 2015). Although the magnitude of repetition suppression BOLD effects can be correlated to behavior, particularly in priming or memory paradigms (Dobbins et al., 2004; Wig et al., 2005; Voss et al., 2009), we did not observe any meaningful brain-behavior relationships. The challenges associated with establishing brain-behavior correlations with fMRI data are well-documented (Rousselet and Pernet, 2012) and the absence of such correlations in our study may be due to a number of factors including the nature of our behavioral paradigm which was primarily designed to verify participants’ compliance on the frequency monitoring task rather than to provide robust performance estimates or to quantify perceptual sensitivities. Alternative behavioral paradigms and fMRI designs are likely required to establish the relationship between BOLD signal changes and temporal frequency perception.

Our within-modality adaptation results may reflect population-level frequency tuning for auditory and tactile stimulation in both parietal and temporal cortex. For auditory regions, the observation of auditory and tactile frequency adaptation effects is perhaps unsurprising given the established existence of neural populations tuned for frequency (Bendor and Wang, 2005; Wang and Walker, 2012) and the common observation that auditory cortical regions can respond to tactile stimulation alone (Foxe et al., 2002; Kayser et al., 2005; Schurmann et al., 2006; Nordmark et al., 2012; Perez-Bellido et al., 2017), even at the unit level (Fu et al., 2003; Lemus et al., 2010). Our data also revealed repetition suppression effects for both modalities in somatosensory areas. The general observation of auditory responses in parietal regions is consistent with other fMRI studies (Beauchamp and Ro, 2008; Liang et al., 2013; Perez-Bellido et al., 2017). Furthermore, causal manipulation of parietal cortex activity can selectively disrupt auditory frequency perception (Convento et al., 2018). However, the inference that neurons in somatosensory regions are tuned for frequency is more tenuous because there has been scant neurophysiological evidence for explicit rate-based frequency tuning in somatosensory neurons unlike their counterparts in the auditory system. Indeed, despite the suggestion that explicit coding for vibration frequencies exists based on modeling studies (Bensmaia et al., 2005; Crommett et al., 2017; Rahman and Yau, 2019), only frequencies in the flutter range (<65Hz) appear to be encoded in spike rates (Salinas et al., 2000; Saal et al., 2016); higher frequencies (>100Hz) like those tested here only appear to be represented in a spike timing code in primate somatosensory cortex (Mountcastle et al., 1969; Ferrington and Rowe, 1980; Harvey et al., 2013). Note that a recent study reported explicit coding for high frequency vibrations in somatosensory neurons for the first time in the mouse (Prsa et al., 2019). Thus, while our results imply that frequency-tuned neural populations may reside in parietal cortex of primates, this conjecture remains to be tested in neurophysiology experiments.

The recruitment of auditory cortex by touch has been interpreted as evidence for a function-based organization scheme in which putative auditory neural circuits that are specialized to represent temporal frequency information also support tactile frequency processing (Yau et al., 2009b). Supramodal processing systems have been invoked for the processing of shape (Pascual-Leone and Hamilton, 2001; Lacey et al., 2009) and motion (Konkle et al., 2009; Wacker et al., 2011). Despite our finding that auditory and tactile frequency-selective responses overlap at a voxel level in many regions, our collective results would appear to argue against a strong supramodal hypothesis for a number of reasons. First, if the same neural populations represented auditory and somatosensory frequency, we may have expected that the magnitude of adaptation effects would be positively correlated between the two modalities. We instead only observed negative or weak correlations over all ROIs or in individual ROIs. Second, if somatosensory and auditory frequency information were carried in overlapping populations, we may have expected to find repetition suppression effects with the crossmodal events that were comparable to those seen with unisensory events. Despite evidence for crossmodal adaptation effects on frequency perception (Levitan et al., 2015; Badde et al., 2016; Crommett et al., 2017) and crossmodal adaptation of fMRI responses with other stimulus features (Tal and Amedi, 2009; Doehrmann et al., 2010), we did not observe significant simple across-modality adaptation effects. With the TA events, significant frequency-dependent patterns were found in anterior parietal cortex and lateral parietal cortex. These patterns were largely explained as a linear combination of 1) frequency-specific excitatory response modulation attributed to the tactile adaptors 2), frequency-specific suppressive response modulation attributed to the auditory probes, and 3) a response component consistent with the activations associated with the TT events. Importantly, each of these response components reflected frequency-dependent processing, so the TA results provide some evidence for the existence of neural circuitry in parietal cortex that responds selectively to auditory and tactile frequency. In sum, while our collective univariate analysis results imply that a number of parietal and temporal regions exhibit frequency-specific responses to auditory and tactile stimulation, these representations may be carried in separate or minimally overlapping neural populations at the voxel level (Driver and Noesselt, 2008; Klemen and Chambers, 2012).

Although exposure to auditory tones has been shown to modulate the subsequent perception of tactile frequency in a frequency-specific manner (Crommett et al., 2017), our fMRI data did not reveal significant response modulation with the AT adaptation events. One possibility is that the response components associated with the tactile probes were too weak relative to the responses associated with the auditory adaptors. Indeed, responses to tactile stimulation tended to be weaker than responses to auditory stimulation (Fig. 4). Likewise, spatial activity patterns were more consistent and robust for auditory stimuli compared to tactile stimuli in the higher-order regions that exhibited prominent multimodal responsiveness (Fig. 8). A strong activity imbalance favoring the auditory modality may have obscured potential frequency-dependent tactile responses or interaction effects. Another possibility is that longer adaptation periods are required to reveal the fMRI correlates of AT adaptation effects. The event-related design used here was only suited to reveal adaptation effects that could emerge within a period of seconds, but perceptual crossmodal adaptation effects have been seen following prolonged adaptation on the order of minutes. Future studies should manipulate the relative strength of the auditory and tactile responses, perhaps by testing auditory band-passed noise stimuli which also interact with touch in a frequency-dependent manner (Yau et al., 2009b; Crommett et al., 2017), and adaptation durations to establish a more comprehensive understanding of audiotactile crossmodal adaptation effects.

While fMRI adaptation has been used extensively to infer population-level tuning properties, there are a number of caveats to consider. First, given the relatively coarse temporal resolution of fMRI and the sluggishness of the BOLD signal, there is ambiguity in relating repetition suppression effects to neural processing in a particular region. As with all fMRI, activation patterns likely reflect the processing of feedback or recurrent signals as much as feedforward responses. Furthermore, adaptation patterns can be inherited from upstream processes (Tolias et al., 2005; Mur et al., 2010). Accordingly, adaptation effects observed in higher-order brain regions may simply reflect the repetition suppression effects in earlier cortical regions or even adaptation in the periphery or subcortical systems. Analogous experiments using imaging and recording methods with higher temporal resolutions will be needed to address these concerns. Similarly, additional experiments that more finely sample the stimulus parameter space (adaptation durations and frequencies) may better enable efforts to link the BOLD adaptation results to neural computations using quantitative encoding models as has been done in studies of compressive temporal summation in the human visual system (Zhou et al., 2017). A more complete sampling of stimulus space may also support pattern component modeling efforts (Diedrichsen et al., 2018) that could link the RSA results to neural encoding models. Such experiments are required to move beyond the more descriptive adaptation results presented here that may only be suited to inform questions regarding the sensory modality preferences of parietal and temporal cortex.

Our multivariate analysis of the task-based data revealed an ROI landscape that was consistent with the traditional view of parietal and temporal cortical systems. Even as we observed multimodal responses in the parietal and temporal regions, there was a general organization seemingly defined according to sensory modality preferences. Crucially, our resting state analyses revealed a highly similar organization pattern between the parietal and temporal regions (Fig. 9). This correspondence was critical because it implies that the ROI landscape cannot be attributed to stimulus-related activity only. In control experiments, we took particular care to confirm that the tactile stimulation used in the scanner were inaudible (Materials and Methods) and that parietal responses to auditory stimulation could not be trivially attributed to mechanical vibrations generated via the in-ear bud headphones (Perez-Bellido et al., 2017). These control experiments alleviate concerns that the overlap between the auditory and tactile responses could be explained by some physical explanation related to poor stimulus control; the resting state results, free of any sensory stimulation effects, directly address this issue. The relationship between intrinsic fluctuations and representational similarity analyses has received recent attention (Henriksson et al., 2015) and our data are consistent with the notion that spatially-patterned intrinsic cortical dynamics could underlie or enhance apparent relationships in the representational geometries over different brain regions. Regardless of how the RSA and functional connectivity results relate, it is indisputable that both reveal a ROI landscape resembling the traditional model of sensory cortex organization. This landscape is also evident in anatomical connectivity profiles determined from invasive tracer studies (Cappe and Barone, 2005; Hackett et al., 2007; Cappe et al., 2009) and diffusion-tensor imaging (Ro et al., 2013). How information is dynamically routed through these connected systems remains an open question. Attention may play a critical role in gating signal transmission between the parietal and temporal networks (Convento et al., 2018). Future studies are required to investigate the state-dependence of frequency information processing in parietal and temporal cortex.

In sum, our results provide evidence for frequency-selective processing of tactile and auditory stimulation over brain regions spanning parietal and temporal cortex. The frequency-selectivity is revealed in unimodal repetition suppression effects as well as the frequency-dependent suppression seen with the auditory probes in the TA events. These results contribute to the growing literature showing the multimodal nature of sensory regions that are traditionally thought to be dedicated to a single modality. Our data highlights the feature-specificity of the multimodal responses. Importantly, we also see evidence of distinct sensory systems organized according to their preferences to audition or touch. Thus, our results reveal that multimodal frequency responses can be distributed over traditionally-defined sensory cortical systems. Because we only characterized responses to sinusoidal stimulation, our analyses could only address processing differences associated with frequency or modality manipulations. Future studies will need to test more complex and naturalistic stimuli, like frequency sweeps (Crommett et al., 2018) or textures (Lederman, 1979; Yau et al., 2009a; Manfredi et al., 2014), to more thoroughly investigate how auditory and tactile representations are elaborated and maintained over hierarchical processing streams in parietal and temporal cortex.

## Acknowledgements

We would like to thank Yau Lab members for helpful discussions and J. Holden for technical assistance with the CM3. This work was supported by R01NS097462 (JMY), Alfred P. Sloan Research Fellowship (JMY), support from the Caroline Wiess Law Fund for Research in Molecular Medicine (JMY), and the Office of Naval Research (MT). We would like to thank Krista Runge and BCM’s Core for Advanced MRI (CAMRI) for technical support.

## References

Alais D, Orchard-Mills E, Van der Burg E (2015) Auditory frequency perception adapts rapidly to the immediate past. Atten Percept Psychophys 77:896–906.

Badde S, Thomaschewski L, Stoffregen H, Roder B (2016) Adapting to visual and auditory low frequency modulated stimuli induced enhanced tactile frequency discrimination. Soc Neurosci.

Barnes KA, Anderson KM, Plitt M, Martin A (2014) Individual differences in intrinsic brain connectivity predict decision strategy. J Neurophysiol 112:1838–1848.

Barron HC, Garvert MM, Behrens TEJ (2016) Repetition suppression: a means to index neural representations using BOLD? Philos Trans R Soc B Biol Sci 371:20150355.

Beauchamp MS, Ro T (2008) Neural Substrates of Sound-Touch Synesthesia after a Thalamic Lesion. J Neurosci 28:13696–13702.

Bendor D, Wang X (2005) The neuronal representation of pitch in primate auditory cortex. Nature 436:1161–1165.

Bendor D, Wang X (2007) Differential neural coding of acoustic flutter within primate auditory cortex. Nat Neurosci 10:763–771.

Bensmaia S, Hollins M, Yau J (2005) Vibrotactile intensity and frequency information in the pacinian system: a psychophysical model. Percept Psychophys 67:828–841.

Cappe C, Barone P (2005) Heteromodal connections supporting multisensory integration at low levels of cortical processing in the monkey. Eur J Neurosci 22:2886–2902.

Cappe C, Morel A, Barone P, Rouiller EM (2009) The thalamocortical projection systems in primate: an anatomical support for multisensory and sensorimotor interplay. Cereb Cortex 19:2025–2037.

Convento S, Rahman MS, Yau JM (2018) Selective Attention Gates the Interactive Crosssmodal Coupling between Perceptual Systems. Curr Biol 28:746–752.

Convento S, Wegner-Clemens KA, Yau JM (2019) Reciprocal interactions between audition and touch in flutter frequency perception. Multisens Res 32:67–85.

Cox RW (1996) AFNI: software for analysis and visualization of functional magnetic resonance neuroimages. Comput Biomed Res 29:162–173.

Crommett LE, Madala D, Yau JM, Crommett LE, Yau JM (2018) Multisensory Perceptual Interactions Between Higher-Order Temporal Frequency Signals. J Exp Psychol Gen.

Crommett LE, Perez-Bellido A, Yau JM (2017) Auditory adaptation improves tactile frequency perception. J Neurophysiol:jn 00783 2016.

Destrieux C, Fischl B, Dale A, Halgren E (2010) Automatic parcellation of human cortical gyri and sulci using standard anatomical nomenclature. Neuroimage 53:1–15.

Diedrichsen J, Yokoi A, Arbuckle SA (2018) Pattern component modeling: A flexible approach for understanding the representational structure of brain activity patterns. Neuroimage 180:119–133.

Dobbins IG, Schnyer DM, Verfaellie M, Schacter DL (2004) Cortical activity reductions during repetition priming can result from rapid response learning. Nature 428:316–319.

Doehrmann O, Weigelt S, Altmann CF, Kaiser J, Naumer MJ (2010) Audiovisual Functional Magnetic Resonance Imaging Adaptation Reveals Multisensory Integration Effects in Object-Related Sensory Cortices. J Neurosci 30:3370–3379.

Driver J, Noesselt T (2008) Multisensory interplay reveals crossmodal influences on “sensory-specific” brain regions, neural responses, and judgments. Neuron 57:11–23.

Ejaz N, Hamada M, Diedrichsen J (2015) Hand use predicts the structure of representations in sensorimotor cortex. Nat Neurosci 18:1034–1040.

Ferrington DG, Rowe MJ (1980) Differential contributions to coding of cutaneous vibratory information by cortical somatosensory areas I and II. J Neurophysiol 43:310–331.

Fischl B, Sereno MI, Dale AM (1999) Cortical surface-based analysis. II: Inflation, flattening, and a surface-based coordinate system. Neuroimage 9:195–207.

Foxe JJ, Wylie GR, Martinez A, Schroeder CE, Javitt DC, Guilfoyle D, Ritter W, Murray MM (2002) Auditory-somatosensory multisensory processing in auditory association cortex: an fMRI study. J Neurophysiol 88:540–543.

Fu KM, Johnston TA, Shah AS, Arnold L, Smiley J, Hackett TA, Garraghty PE, Schroeder CE (2003) Auditory cortical neurons respond to somatosensory stimulation. J Neurosci 23:7510–7515.

Ghazanfar AA, Schroeder CE (2006) Is neocortex essentially multisensory? Trends Cogn Sci 10:278–285.

Goble AK, Hollins M (1993) Vibrotactile adaptation enhances amplitude discrimination. J Acoust Soc Am 93:418–424.

Goble AK, Hollins M (1994) Vibrotactile adaptation enhances frequency discrimination. J Acoust Soc Am 96:771–780.

Grill-Spector K, Henson R, Martin A (2006) Repetition and the brain: neural models of stimulus-specific effects. Trends Cogn Sci 10:14–23.

Hackett TA, Smiley JF, Ulbert I, Karmos G, Lakatos P, de la Mothe LA, Schroeder CE (2007) Sources of somatosensory input to the caudal belt areas of auditory cortex. Perception 36:1419–1430.

Harvey MA, Saal HP, Dammann 3rd JF, Bensmaia SJ (2013) Multiplexing stimulus information through rate and temporal codes in primate somatosensory cortex. PLoS Biol 11:e1001558.

Hegner YL, Saur R, Veit R, Butts R, Leiberg S, Grodd W, Braun C (2007) BOLD adaptation in vibrotactile stimulation: neuronal networks involved in frequency discrimination. J Neurophysiol 97:264–271.

Henriksson L, Khaligh-Razavi SM, Kay K, Kriegeskorte N (2015) Visual representations are dominated by intrinsic fluctuations correlated between areas. Neuroimage 114:275–286.

Hollins M, Goble AK, Whitsel BL, Tommerdahl M (1990) Time course and action spectrum of vibrotactile adaptation. Somatosens Mot Res 7:205–221.

Jo HJ, Saad ZS, Simmons WK, Milbury LA, Cox RW (2010) Mapping sources of correlation in resting state FMRI, with artifact detection and removal. Neuroimage 52:571–582.

Kayser C, Petkov CI, Augath M, Logothetis NK (2005) Integration of touch and sound in auditory cortex. Neuron 48:373–384.

Klemen J, Chambers CD (2012) Current perspectives and methods in studying neural mechanisms of multisensory interactions. Neurosci Biobehav Rev 36:111–133.

Konkle T, Wang Q, Hayward V, Moore CI (2009) Motion aftereffects transfer between touch and vision. Curr Biol 19:745–750.

Krekelberg B, Boynton GM, van Wezel RJ (2006) Adaptation: from single cells to BOLD signals. Trends Neurosci 29:250–256.

Lacey S, Tal N, Amedi A, Sathian K (2009) A putative model of multisensory object representation. Brain Topogr 21:269–274.

Lattner S, Meyer ME, Friederici AD (2005) Voice perception: Sex, pitch, and the right hemisphere. Hum Brain Mapp 24:11–20.

Lederman SJ (1979) Auditory texture perception. Perception 8:93–103.

Lemus L, Hernandez A, Luna R, Zainos A, Romo R (2010) Do sensory cortices process more than one sensory modality during perceptual judgments? Neuron 67:335–348.

Levitan CA, Ban Y-HA, Stiles NRB, Shimojo S (2015) Rate perception adapts across the senses: evidence for a unified timing mechanism. Sci Rep 5:8857.

Liang M, Mouraux A, Hu L, Iannetti GD (2013) Primary sensory cortices contain distinguishable spatial patterns of activity for each sense. Nat Commun 4:1979.

Manfredi LR, Saal HP, Brown KJ, Zielinski MC, Dammann 3rd JF, Polashock VS, Bensmaia SJ (2014) Natural scenes in tactile texture. J Neurophysiol 111:1792–1802.

Mazziotta J et al. (2001) A probabilistic atlas and reference system for the human brain: International Consortium for Brain Mapping (ICBM). Philos Trans R Soc L B Biol Sci 356:1293–1322.

Miller LE, Montroni L, Koun E, Salemme R, Hayward V, Farnè A (2018) Sensing with tools extends somatosensory processing beyond the body. Nature 561:239–242.

Millin R, Kolodny T, Flevaris A V, Kale AM, Schallmo M-P, Bernier RA, Murray SO (2018) Disrupted neural adaptation in autism spectrum disorder. Elife 7:1–15.

Mountcastle VB, Talbot WH, Sakata H, Hyvärinen J (1969) Cortical neuronal mechanisms in flutter-vibration studied in unanesthetized monkeys. Neuronal periodicity and frequency discrimination. J Neurophysiol 32:452–484.

Mur M, Ruff DA, Bodurka J, Bandettini PA, Kriegeskorte N (2010) Face-identity change activation outside the face system: “release from adaptation” may not always indicate neuronal selectivity. Cereb Cortex 20:2027–2042.

Nordmark PF, Pruszynski JA, Johansson RS (2012) BOLD responses to tactile stimuli in visual and auditory cortex depend on the frequency content of stimulation. J Cogn Neurosci 24:2120–2134.

Oldfield RC (1971) The assessment and analysis of handedness: the Edinburgh inventory. Neuropsychologia 9:97–113.

Parra LC, Pearlmutter BA (2007) Illusory percepts from auditory adaptation. J Acoust Soc Am 121:1632–1641.

Pascual-Leone A, Hamilton R (2001) The metamodal organization of the brain. Prog Brain Res 134:427–445.

Perez-Bellido A, Barnes KA, Crommett LE, Yau JM (2017) Auditory frequency representations in human somatosensory cortex. Cereb Cortex.

Prsa M, Morandell K, Cuenu G, Huber D (2019) Feature-selective encoding of substrate vibrations in the forelimb somatosensory cortex. Nature 567:384–388.

Rahman MS, Yau JM (2019) Somatosensory interactions reveal feature-dependent computations. J Neurophysiol.

Ro T, Ellmore TM, Beauchamp MS (2013) A neural link between feeling and hearing. Cereb Cortex 23:1724–1730.

Romo R, Salinas E (2003) Flutter discrimination: neural codes, perception, memory and decision making. Nat Rev Neurosci 4:203–218.

Rousselet GA, Pernet CR (2012) Improving standards in brain-behavior correlation analyses. Front Hum Neurosci 6.

Saad ZS, Reynolds RC (2012) SUMA. Neuroimage 62:768–773.

Saal HP, Wang X, Bensmaia SJ (2016) Importance of spike timing in touch: an analogy with hearing? Curr Opin Neurobiol 40:142–149.

Salinas E, Hernandez A, Zainos A, Romo R (2000) Periodicity and firing rate as candidate neural codes for the frequency of vibrotactile stimuli. J Neurosci 20:5503–5515.

Schurmann M, Caetano G, Hlushchuk Y, Jousmaki V, Hari R (2006) Touch activates human auditory cortex. Neuroimage 30:1325–1331.

Solomon SG, Kohn A (2014) Moving sensory adaptation beyond suppressive effects in single neurons. Curr Biol 24:R1012–22.

Tal N, Amedi A (2009) Multisensory visual-tactile object related network in humans: insights gained using a novel crossmodal adaptation approach. Exp Brain Res 198:165–182.

Tame L, Braun C, Lingnau A, Schwarzbach J, Demarchi G, Li Hegner Y, Farne A, Pavani F (2012) The contribution of primary and secondary somatosensory cortices to the representation of body parts and body sides: an fMRI adaptation study. J Cogn Neurosci 24:2306–2320.

Titze IR (1994) Principles of Voice Production. Prentice Hall.

Tolias AS, Keliris GA, Smirnakis SM, Logothetis NK (2005) Neurons in macaque area V4 acquire directional tuning after adaptation to motion stimuli. Nat Neurosci 8:591–593.

Tommerdahl M, Hester KD, Felix ER, Hollins M, Favorov O V, Quibrera PM, Whitsel BL (2005) Human vibrotactile frequency discriminative capacity after adaptation to 25 Hz or 200 Hz stimulation. Brain Res 1057:1–9.

Vergara J, Rivera N, Rossi-Pool R, Romo R (2016) A Neural Parametric Code for Storing Information of More than One Sensory Modality in Working Memory. Neuron 89:54–62.

Voss JL, Hauner KKY, Paller KA (2009) Establishing a relationship between activity reduction in human perirhinal cortex and priming. Hippocampus 19:773–778.

Wacker E, Spitzer B, Lutzkendorf R, Bernarding J, Blankenburg F (2011) Tactile motion and pattern processing assessed with high-field FMRI. PLoS One 6:e24860.

Wang X, Walker KMM (2012) Neural Mechanisms for the Abstraction and Use of Pitch Information in Auditory Cortex. J Neurosci 32:13339–13342.

Wig GS, Grafton ST, Demos KE, Kelley WM (2005) Reductions in neural activity underlie behavioral components of repetition priming. Nat Neurosci 8:1228–1233.

Yau JM, Hollins M, Bensmaia SJ (2009a) Textural timbre: The perception of surface microtexture depends in part on multimodal spectral cues. Commun Integr Biol 2:344–346.

Yau JM, Olenczak JB, Dammann JF, Bensmaia SJ (2009b) Temporal Frequency Channels Are Linked across Audition and Touch. Curr Biol 19:561–566.

Yau JM, Weber AI, Bensmaia SJ (2010) Separate mechanisms for audio-tactile pitch and loudness interactions. Front Psychol 1:doi: 10.3389/fpsyg.2010.00160.

Zhou J, Benson NC, Kay KN, Winawer J (2017) Compressive Temporal Summation in Human Visual Cortex. J Neurosci 38:691–709.

Zwicker E (1964) Negative afterimage in hearing. J Acoust Soc Am 36:2413–2415.

